# A primary rodent triculture model to investigate the role of glia-neuron crosstalk in regulation of neuronal activity

**DOI:** 10.1101/2022.09.30.510271

**Authors:** Leela Phadke, Dawn H. W. Lau, Nozie D. Aghaizu, Shania Ibarra, Carmen M. Navarron, Lucy Granat, Lorenza Magno, Paul Whiting, Sarah Jolly

## Abstract

Neuroinflammation and hyperexcitability have been implicated in the pathogenesis of neurodegenerative disease, and new models are required to investigate the cellular crosstalk involved in these processes. We developed an approach to generate a quantitative and reproducible triculture system that is suitable for pharmacological studies. While primary rodent cells were previously grown in a coculture medium formulated to support only neurons and astrocytes, we now optimised a protocol to generate tricultures containing neurons, astrocytes and microglia by culturing in a medium designed to support all three cell types and adding exogenous microglia to cocultures. Immunocytochemistry was used to confirm the intended cell types were present. The percentage of ramified microglia in the tricultures decreases as the number of microglia present increases. Multi-electrode array (MEA) recordings indicate that microglia in the triculture model suppress neuronal activity in a dose-dependent manner. Neurons in both cocultures and tricultures were responsive to the potassium channel blocker 4-aminopyridine (4-AP) suggesting that neurons remained viable and functional in the triculture model. Furthermore, suppressed neuronal activity in tricultures correlated with decreased densities of dendritic spines and of the postsynaptic protein Homer1 along dendrites, indicative of a direct or indirect effect of microglia on synapse function.

We thus present a functional triculture model, which, due to its more complete cellular composition, is a more relevant model than standard cocultures. The model can be used to probe glia-neuron interactions and subsequently aid the development of assays for drug discovery, using neuronal excitability as a functional endpoint.

## 1. Introduction

Alzheimer’s Disease (AD) is the most common cause of dementia, accounting for an estimated 60–80% of cases worldwide (Alzheimer’s Association, 2020). The two main pathological hallmarks of AD in the brain are amyloid-β (Aβ) plaques and abnormal tau tangles, and it has been proposed that Aβ accumulation initiates a cascade of events that ultimately results in neuronal damage. However, the linearity of this cascade remains controversial and accumulating evidence suggests that many processes are involved in the pathogenesis of AD (De Strooper and Karran, 2016). Neuronal hyperexcitability has been implicated in the pathogenesis of neurodegenerative disease and correlates with cognitive decline in patients with AD (Vossel et al., 2013; Harris et al., 2020). In addition, studies have found that elevated levels of inflammatory markers are present in patients with AD (Brosseron et al., 2018), and a number of AD risk genes are associated with innate immune functions (Efthymiou and Goate, 2017; Bellenguez et al., 2022). Therefore, neuroinflammation is thought to play a significant role in the pathogenesis of the disease and the immune cells involved in this process are microglia and astrocytes.

Astrocytes are the most abundant cell type in the central nervous system (CNS). Their functions include the maintenance of neurotransmitter homeostasis, synapse formation, metabolic and neurotrophic support for neurons (Rouach et al., 2008; Eroglu and Barres, 2010). Astrocytes are known to respond to pathological insults through reactive gliosis, which involves morphological, molecular, and functional remodelling and they can also become atrophic, showing reduced volume and a reduced number of processes (Arranz and De Strooper, 2019; Escartin et al., 2021). Microglia are the resident macrophages of the CNS that play important roles in synaptic pruning, neuronal apoptosis and maintenance of synaptic plasticity (Schafer et al., 2012; Weinhard et al., 2018; York et al., 2018). Microglia have also been shown to respond to neuronal activation by suppressing neuronal activity (Badimon et al., 2020). Homeostatic, ramified microglia constantly survey the CNS with their processes, sensing damage signals and mediating a response towards the site of injury (Nimmerjahn et al., 2005). Microglia *in vivo* are highly dynamic and heterogeneous, allowing them to achieve a range of responses depending on the environmental context. It remains unclear whether microglial function in neurodegenerative disease is beneficial but insufficient, or whether they are effective in early disease before losing their efficacy or becoming detrimental (Masuda et al., 2020).

Analysis of tissue from the mouse CNS during homeostasis reveals specific time- and region-dependent subtypes of microglia, and additional context-dependent subtypes are found in mouse models of neurodegeneration and demyelination. These subtypes correspond to similar clusters identified in healthy human brains and the brains of multiple sclerosis patients (Masuda et al., 2019). This heterogeneity is difficult to capture in a classically used monoculture system. Developing more complex cultures with other cell types combined in the same environment may provide a better model of the diverse phenotypic states observed *in vivo*. Neurons, astrocytes and microglia act in a synchronized manner, and communication between all these cell types can be disrupted by, and contribute to, neurodegenerative diseases. For example, in the presence of Aβ, microglia become responsive to IL-3 produced by astrocytes and undergo transcriptional and phenotypic changes leading them to a more reactivate state that can restrict AD pathology (McAlpine et al., 2021). Aβ was also shown to activate the NF-κB pathway in astrocytes and increase the release of complement C3, resulting in neuronal dysfunction and microglial activation via C3a receptor signalling (Lian et al., 2015). Conversely, activated microglia induce a specific state of reactive astrocytes by secreting Il-1α, TNF and C1q and this population of astrocytes has been shown to be neurotoxic (Liddelow et al., 2017; Guttenplan et al., 2021). Furthermore, selective blocking of Aβ-induced microglial activation via GLP-1R activation inhibits formation of reactive astrocytes and preserves neuronal viability in AD models (Park et al., 2021). Dysregulated neuronal–glial crosstalk has also been observed in human AD. Under physiological conditions, neuronal–microglial crosstalk via CD200–CD200R and CX3CL1–CX3CR1 signalling is thought to regulate the homeostatic function of microglia (Simon et al., 2019). In the brains of AD patients, however, expression of CD200, CD200R and CX3CR1 is reduced, suggesting a loss of physiological constraints on microglial function (Walker et al., 2009).

There is a need for more complex *in vitro* systems able to model the crosstalk between different cell types in the CNS. Cocultures of neurons and astrocytes are an established method of studying neuroinflammation *in vitro* but these models are not able to capture the interactions with microglia or how both astrocytes and microglia may affect neuronal activity. We thus generated an experimental rodent triculture platform comprised of all three major cell types associated with neuroinflammation (neurons, astrocytes, and microglia) that is quantitative and reproducible to allow for pharmacological studies and screening. Primary rodent cells were maintained in a serum-free culture media that we optimised to support all three cell types. To validate our platform, we utilised immunostaining followed by High Content Imaging System and Multi-Electrode Array (MEA) as endpoints. MEA technology is a powerful tool for quantifying the activity of neuronal networks in a fast and non-invasive manner. We showed that this triculture model can be maintained for at least 17 days *in vitro* and that microglia in the model do not affect the number of neurons but suppress neuronal activity in a dose-dependent manner. Here we developed an assay for investigating glia-mediated regulation of cortical neuron activity, which is amenable to pharmacological modulation. This platform will facilitate drug discovery by enabling target identification studies as well as drug screening.

## 2. Results

### 2.1. Generation of a triculture model with rodent primary neurons, astrocytes and microglia

Our aim was to establish a triculture model allowing for the investigation of glial contribution to neuronal activity. Primary cortical cells from E18 rat embryos were cultured in neuron-astrocyte coculture media for 6 days (Fig. 1A). Early experiments showed that when microglia were added to the culture at DIV8 and cultured in coculture media, less than 1% microglia were present by DIV14 (data not shown). TGF-β2, IL-34 and cholesterol have been identified as the factors essential for microglial survival in serum-free, defined medium conditions. Microglia in isolation can be cultured in a serum-free ‘TICH’ media supplemented with TGF-β2, IL-34 and cholesterol as well as heparan sulphate, which improves cell adhesion and process extension (Bohlen et al., 2017). Microglia were initially plated on cocultures in TICH media but experiments showed that this addition of microglia or addition of TICH media alone caused a significant reduction in the number of NeuN-positive neuronal nuclei present (data not shown). To support microglial survival and avoid the neurotoxicity due to the addition of microglia in TICH media, we developed a triculture media supporting all three cell types, which was composed of the coculture media supplemented with TGF-β2, IL-34 and cholesterol.

**Figure 1:**
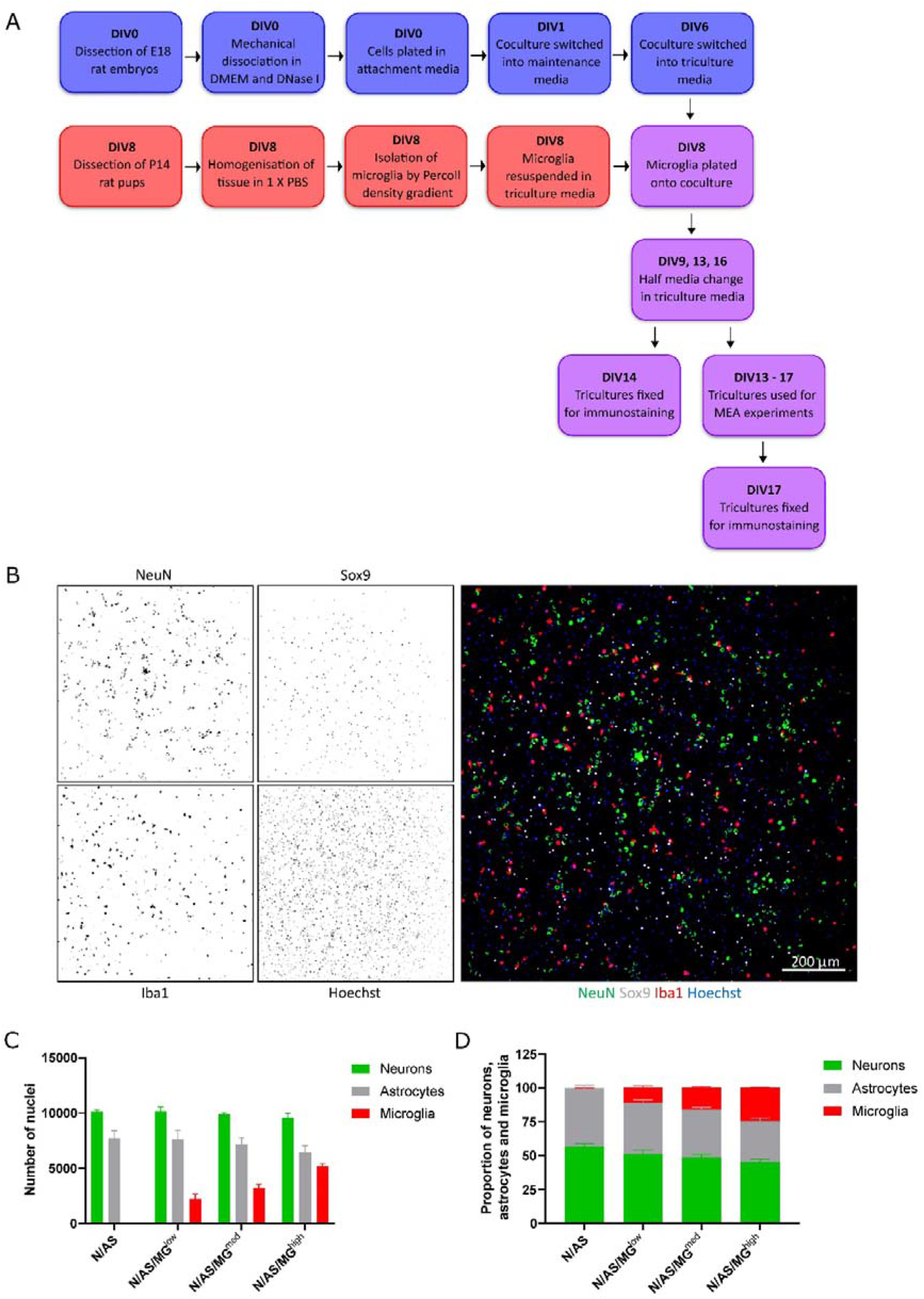
Development of a triculture system with rodent neurons, astrocytes and microglia. **A**. Schematic workflow of neurons, astrocytes and microglia plating, maintenance, and maturation. Blue = coculture of neurons and astrocytes. Red = microglia. Purple = triculture of neurons, astrocytes and microglia. Mature cultures (DIV13+) were used for various experimental treatments and conditions. At the end of the experiments, fixed cells were processed for immunostaining. **B, C, D.** Quantification of cell numbers by immunocytochemistry. (**B**). Representative immunostaining image used for quantification showing NeuN (neuronal marker), Sox9 (astrocyte marker) and Iba1 (microglia marker) expression at DIV14 in triculture N/AS/MG^high^. Merge shows staining of NeuN (green), Sox9 (white), Iba1 (red) and Hoechst (blue). Scale bar = 200 μm. (**C**). Cell count analysis of the total number of cells per well performed on cocultures and tricultures at DIV14 using Harmony software. There was no significant difference in astrocyte populations in the tricultures compared to the coculture (N/AS/MG^low^ *p* > 0.9999, N/AS/MG^med^ *p* = 0.9993, N/AS/MG^high^ *p* = 0.7218, 2-way ANOVA, post-hoc Tukey’s multiple comparisons test). Data shown as mean ± SEM and is representative of 3 separate experiments. (**D**). Cell count analysis shown as proportions of the total number of neurons, astrocytes and microglia. Data shown as mean ± SEM and is representative of 3 independent experiments.

To allow the neurons and astrocytes to adjust to the additional media components prior to addition of the microglia on DIV8, cocultures were switched into triculture media 48 hours before on DIV6. Exogenous microglia isolated from P14 rats were resuspended in triculture media and plated onto the coculture on DIV8 at three different ratios: 1 neuron/astrocyte:0.1 microglia, 1 neuron/astrocyte:0.3 microglia and 1 neuron/astrocyte:1.5 microglia, referred to as N/AS/MG^low^, N/AS/MG^med^ and N/AS/MG^high^ respectively (Fig. 1A). These ratios were chosen to cover the range of cell densities that have been reported in the human brain, as the exact numbers remain unknown and appear to vary between different brain regions (Herculano-Houzel, 2014; von Bartheld et al., 2016). Ionized calcium-binding adapter molecule 1 (Iba1) is a protein expressed specifically by microglia in the brain (Ito et al., 1998). Immunostaining for Iba1 showed that at DIV14 there was a population of microglia present in the triculture which increased dose-dependently and was absent in the coculture (Fig. 1B and 1C). Immunostaining for the neuronal marker NeuN and astrocyte-specific nuclear marker Sox9 indicated a healthy population of neurons and astrocytes in both the cocultures and tricultures (Fig. 1B and 1C). The number of neurons was not affected by the presence of the microglia (Fig. 1C). While the analysis also did not reveal a significant difference in astrocyte populations in the tricultures compared to the coculture (N/AS/MG^low^ *p* > 0.9999, N/AS/MG^med^ *p* = 0.9993, N/AS/MG^high^ *p* = 0.7218), the results suggest that the N/AS/MG^high^ triculture may contain slightly lower numbers of astrocytes than the coculture (three independent biological repeats were quantified). As a proportion of the total number of neurons, astrocytes and microglia in culture, cocultures contained 56.89±1.952% neurons, 43.10±1.952% astrocytes and no microglia. The N/AS/MG^low^ triculture contained 51.13±3.133% neurons, 37.98±2.031% astrocytes and 10.89±1.5665% microglia. The N/AS/MG^med^ triculture contained 48.82±2.279% neurons, 35.32±1.527% astrocytes and 15.89±0.8791% microglia. The N/AS/MG^high^ triculture contained 45.22±2.169% neurons, 30.33±2.192 SE astrocytes and 24.45±0.2614% microglia (Fig 1D). Note that the NeuN staining was sometimes weak and/or did not mark all the neurons. Additional staining for the Oligodendrocyte transcription factor 2 (Olig2) (Valério-Gomes et al., 2018) revealed that both cocultures and tricultures contained 3-8% of cells from the oligodendrocyte lineage (data not shown).

Microglia *in vitro* display a high degree of heterogeneity in terms of morphology. Homeostatic microglia display ramifications and processes, whereas activated microglia are more amoeboid. As microglial morphology gives an indication as to the phenotypic state of the cell, we performed further analysis on the tricultures to classify the morphology of microglia as either ramified or amoeboid. While the difference between the three triculture conditions is not statistically significant (*p* = 0.2440), the trend indicates that the percentage of ramified microglia present in the triculture decreases dose-dependently with increasing numbers of microglia. The difference between the percentage of ramified microglia in the N/AS/MG^low^ triculture and the N/AS/MG^high^ triculture (*p* = 0.2201) is larger than the difference between the N/AS/MG^low^ triculture and the N/AS/MG^med^ triculture (*p* = 0.5839) (Fig 2A and 2B). The average percentage of ramified microglia from three biological repeats was 26.45±4.272% for the N/AS/MG^low^ triculture, 20.53±4.875% for the N/AS/MG^med^ triculture and 15.61±2.668% for the N/AS/MG^high^ triculture (Fig 2B).

**Figure 2:**
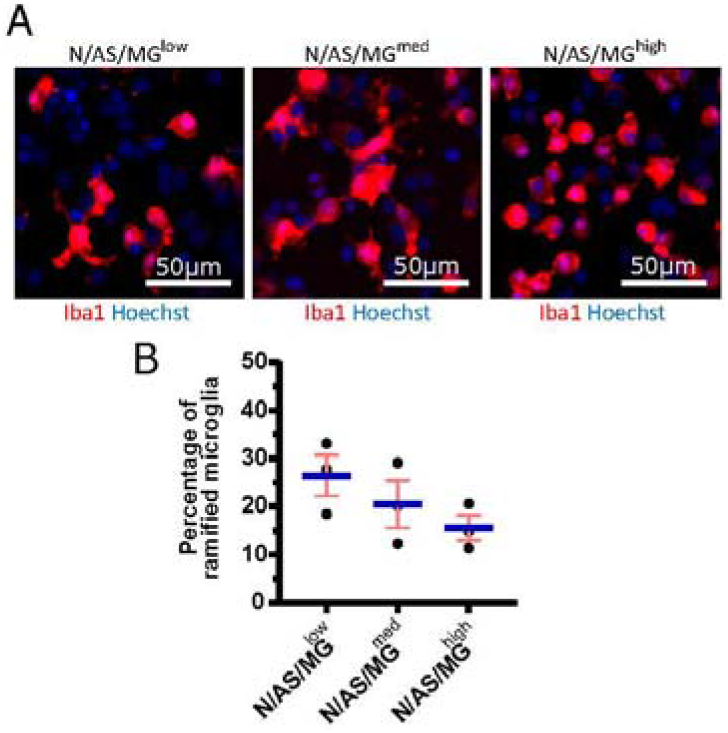
Microglia in tricultures with higher numbers of microglia display a less ramified morphology than tricultures with less microglia. (**A**). Representative immunostaining images used for morphology analysis showing Iba1 expression at DIV14 in tricultures N/AS/MG^low^, N/AS/MG^med^ and N/AS/MG^high^ 1:3. Merge shows staining of Iba1 (red) and Hoechst (blue). Scale bar = 50 μm. (**B**). Morphology analysis performed on Iba1-positive cells in tricultures at DIV14 using Harmony software. The difference between the three triculture conditions is not statistically significant (*p* = 0.2440, one-way ANOVA). However, the difference between the percentage of ramified microglia in the N/AS/MG^low^ triculture and the N/AS/MG^high^ triculture (*p* = 0.2201, one-way ANOVA, post-hoc Tukey’s multiple comparisons test) is larger than the difference between the N/AS/MG^low^ triculture and the N/AS/MG^med^ triculture (*p* = 0.5839, one-way ANOVA, post-hoc Tukey’s multiple comparisons test). Data shown as mean ± SEM and is representative of 3 independent experiments.

### 2.2. Use of MEA to measure neuronal activity shows that microglia suppress spontaneous neuronal activity

As both microglia and astrocytes can regulate neuronal activity in basal conditions as well as in disease, we set up an assay allowing for the assessment of glia-mediated regulation of cortical neuron activity. MEA is a high throughput assay allowing recording of spontaneous neuronal activity over time in a non-invasive way. Following optimisation of the protocol, neuronal cultures, cocultures and tricultures were cultured on 96-well MEA plates to record neuronal activity (see Methods for detailed protocol). Neuronal cultures and cocultures are plated by spot-coating onto the centre of the electrodes. Upon their addition, microglia attach to the whole well, not only the electrodes (Fig 3A and 3B). Therefore, the final ratios of the tricultures in the MEA plates were lower than the same ratios plated onto imaging plates. The N/AS/MG^med^ and N/AS/MG^high^ tricultures were chosen for experimentation on the MEA to increase the likelihood that the final ratio was sufficient to characterise the effect of the microglia on neuronal activity. Immunostaining for NeuN, Sox9 and Iba1 confirmed that the intended cell types were present at DIV17 in the cocultures and tricultures cultured on MEA plates (Fig 3A and 3B). Spontaneous neuronal activity in neuronal cultures, cocultures and tricultures from five independent biological repeats was recorded daily from DIV13 to DIV17. The software identified action potentials or ‘spikes’, clusters of spikes referred to as bursts, and coordinated clusters of spikes across multiple electrodes referred to as network bursts. The network burst frequency significantly increased from DIV13 to DIV17 in neuronal cultures (Fig 4C, *p* = 0.0483), with the mean firing rate and burst frequency displaying a similar albeit not significant trend. The mean firing rate, burst frequency and network burst frequency significantly increased from DIV13 to DIV17 in the cocultures and both tricultures (Fig 4A, 4B and 4C, Mean firing rate *p* < 0.0001, Burst frequency *p* < 0.0001, Network burst frequency *p* < 0.0001). The mean firing rate was significantly lower in neuronal cultures compared to cocultures at all time points recorded after DIV13 (Fig 4A, DIV14 to DIV17 *p* < 0.0001). Burst frequency was significantly lower in neuronal cultures compared to cocultures at all time points recorded after DIV13 (Fig 4A, DIV14 *p* = 0.002, DIV15 p = 0.0002, DIV16 to DIV17 *p* < 0.0001). Network burst frequency was significantly lower in neuronal cultures compared to cocultures at all time points recorded after DIV14 (Fig 4A, DIV15 p = 0.0318, DIV16 to DIV17 *p* < 0.0001). Microglia suppressed spontaneous neuronal activity in a dose-dependent manner. The mean firing rate was significantly lower in triculture N/AS/MG^high^ compared to coculture N/AS at all time points recorded (Fig 4A, DIV13 *p* = 0.0456, DIV14 to DIV17 *p* < 0.0001). The mean firing rate was significantly lower in triculture N/AS/MG^med^ compared to coculture N/AS at all time points recorded after DIV14 (Fig 4A, DIV15 *p* = 0.0024, DIV16 *p* < 0.0001, DIV17 *p* = 0.0048). Burst frequency was significantly lower in triculture N/AS/MG^high^ compared to coculture N/AS at all time points recorded after DIV13 (Fig 4B, DIV14 *p* = 0.003, DIV15 to DIV17 *p* < 0.0001). Burst frequency was significantly lower in triculture N/AS/MG^med^ compared to coculture N/AS at all time points recorded after DIV13 (Fig 4B, DIV14 *p* = 0.0023, DIV15 *p* = 0.0001, DIV16 to DIV17 *p* < 0.0001). Network burst frequency was significantly lower in triculture N/AS/MG^high^ compared to coculture N/AS at all time points recorded after DIV13 (Fig 4C, DIV14 *p* = 0.0058, DIV15 *p* = 0.0001, DIV16 to DIV17 *p* < 0.0001). Network burst frequency was significantly lower in triculture N/AS/MG^med^ compared to coculture N/AS at all time points recorded between DIV14 and DIV16 (Fig 4C, DIV14 *p* = 0.0112, DIV15 *p* = 0.0007, DIV16 *p* = 0.0004). Representative activity traces (raster plots) at DIV17 from coculture and tricultures show the dose-dependent inhibition of neuronal activity by microglia (Fig 4D, 4E, 4F).

**Figure 3:**
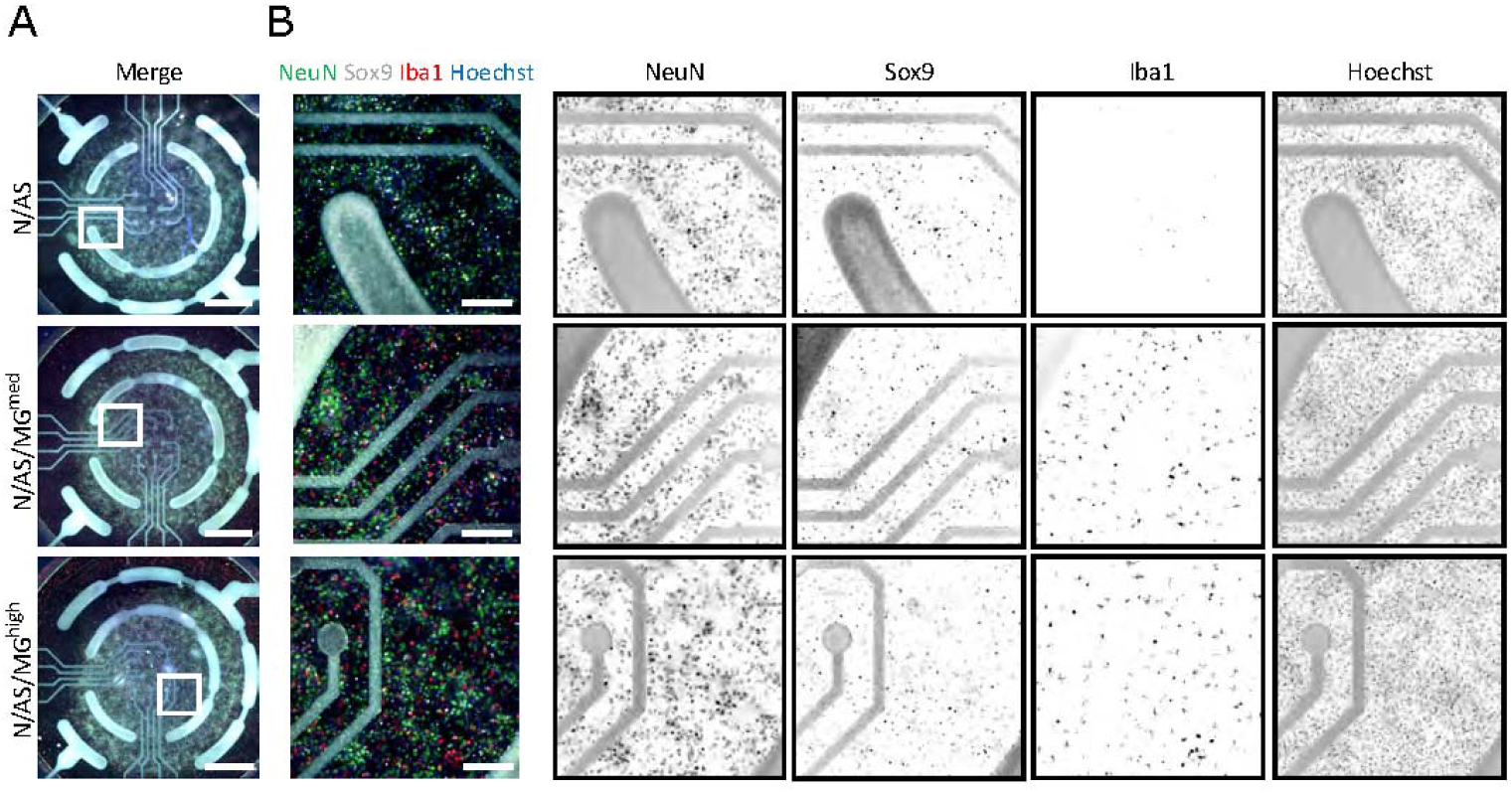
Cocultures and tricultures of rodent cells are cultured on MEA plates to record neuronal activity. (**A**). Representative whole-well immunostaining images showing NeuN, Sox9 and Iba1 expression in cocultures and tricultures used for spontaneous activity recordings in MEA plates at DIV17. Merge shows staining of NeuN (green), Sox9 (white), Iba1 (red) and Hoechst (blue). Scale bar = 1000 μm. White frame = field of view of image (**B**). Representative snapshot immunostaining images showing NeuN, Sox9 and Iba1 expression in cocultures and tricultures used for spontaneous activity recording in MEA plates. Merge shows staining of NeuN (green), Sox9 (white), Iba1 (red) and Hoechst (blue). Scale bar = 200 μm.

**Figure 4:**
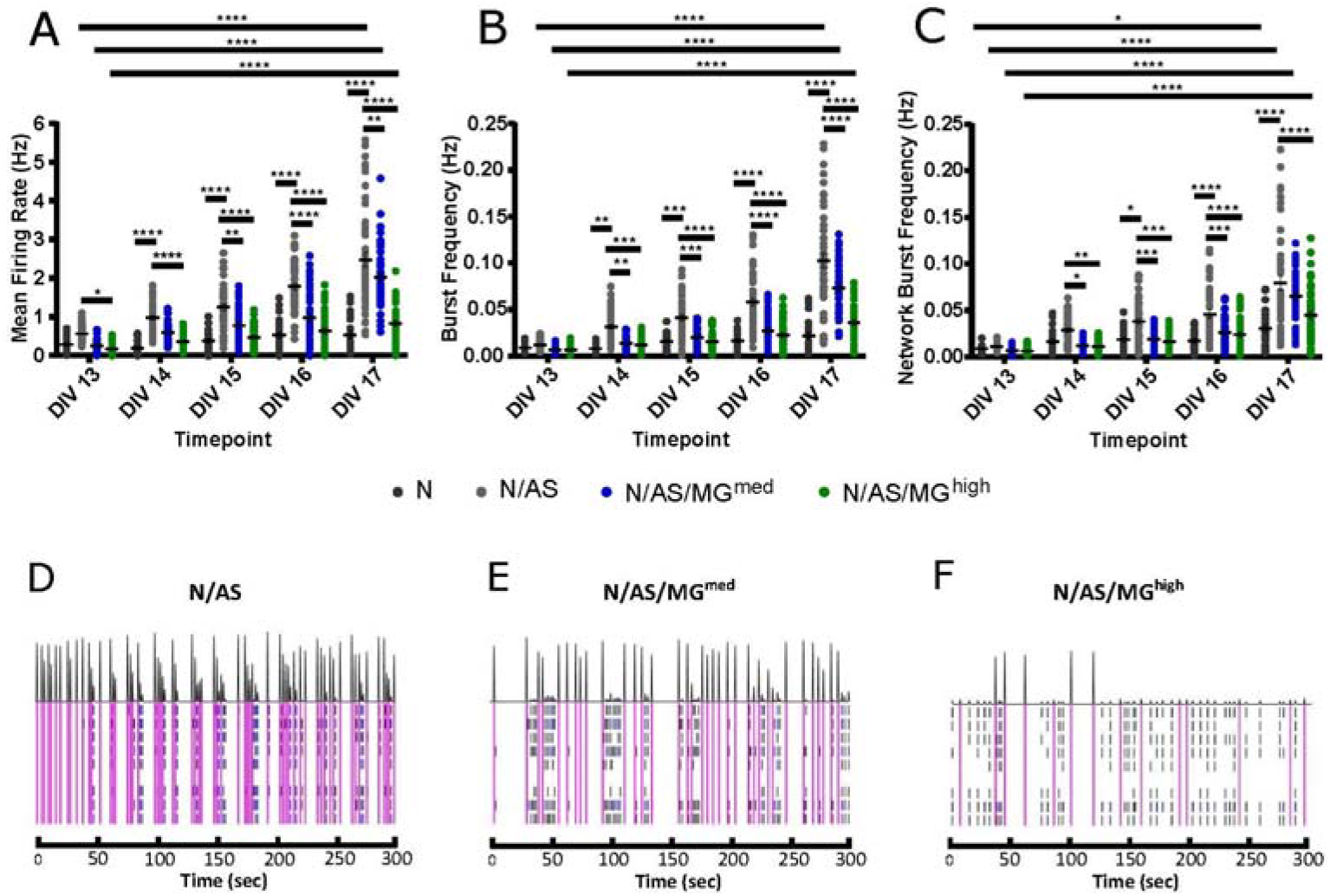
Microglia suppress spontaneous neuronal activity in a dose-dependent manner. (**A-C**). Spontaneous neuronal activity in neuronal cultures, cocultures and tricultures was recorded daily from DIV13 to DIV17. (**A**). Mean firing rate was higher in cocultures compared to neuronal cultures at all time points recorded after DIV13 (Fig 4A, DIV14 to DIV17 *p* < 0.0001, 2-way ANOVA, post-hoc Tukey’s multiple comparisons test). Mean firing rate was higher in cocultures compared to triculture N/AS/MG^high^ at all time points recorded (DIV13 *p* = 0.0456, DIV14 to DIV17 *p* < 0.0001, 2-way ANOVA, post-hoc Tukey’s multiple comparisons test). Mean firing rate was higher in cocultures compared to triculture N/AS/MG^med^ at all time points recorded after DIV14 (Fig 4A, DIV15 *p* = 0.0024, DIV16 *p* < 0.0001, DIV17 *p* = 0.0048, 2-way ANOVA, post-hoc Tukey’s multiple comparisons test). Mean firing rate was higher at DIV17 compared to DIV13 in both the cocultures and tricultures (*p* < 0.0001, 2-way ANOVA, post-hoc Tukey’s multiple comparisons test). (**B**). Burst frequency was higher in cocultures compared to neuronal cultures at all time points recorded after DIV13 (Fig 4A, DIV14 *p* =0.002, DIV15 p = 0.0002, DIV16 to DIV17 *p* < 0.0001, 2-way ANOVA, post-hoc Tukey’s multiple comparisons test). Burst frequency was higher in cocultures compared to triculture N/AS/MG^high^ at all time points recorded after DIV13 (DIV14 *p* = 0.003, DIV15 to DIV17 *p* < 0.0001, 2-way ANOVA, post-hoc Tukey’s multiple comparisons test). Burst frequency was higher in cocultures compared to triculture N/AS/MG^med^ at all time points recorded after DIV13 (DIV14 p = 0.0023, DIV15 p = 0.0001, DIV16 to DIV17 p < 0.0001, 2-way ANOVA, post-hoc Tukey’s multiple comparisons test). Burst frequency was higher at DIV17 compared to DIV13 in both the cocultures and tricultures (*p* < 0.0001, 2-way ANOVA, post-hoc Tukey’s multiple comparisons test). (**C**). Network burst frequency was higher in cocultures compared to neuronal cultures at all time points recorded after DIV14 (Fig 4A, DIV15 p = 0.0318, DIV16 to DIV17 p < 0.0001, 2-way ANOVA, post-hoc Tukey’s multiple comparisons test). Network burst frequency was higher in cocultures compared to triculture N/AS/MG^high^ at all time points recorded after DIV13 (DIV14 *p* = 0.0058, DIV15 *p* = 0.0001, DIV16 *p* < 0.0001, DIV17 *p* < 0.0001, 2-way ANOVA, post-hoc Tukey’s multiple comparisons test). Network burst frequency was higher in cocultures compared to triculture N/AS/MG^med^ at all time points recorded between DIV14 and DIV16 (DIV14 p = 0.0112, DIV15 p = 0.0007, DIV16 p = 0.0004, 2-way ANOVA, post-hoc Tukey’s multiple comparisons test). Network burst frequency was higher at DIV17 compared to DIV13 in both the cocultures and tricultures (*p* < 0.0001, 2-way ANOVA, post-hoc Tukey’s multiple comparisons test). Network burst frequency was also higher at DIV17 compared to DIV13 in neuronal cultures (Fig 4C, p = 0.0483, 2-way ANOVA, post-hoc Tukey’s multiple comparisons test). (**D, E, F**). Representative activity traces at DIV17 from coculture N/AS and tricultures N/AS/MG^med^ and N/AS/MG^high^. Black spikes at the top represent network activity while activity over time recorded on each of the 8 electrodes is shown under. Blue = bursts. Pink = network bursts. Data shown as mean ± SEM and is representative of 5 independent experiments.

### 2.3. Neurons in triculture retain activity capacity despite microglial attenuation of electrophysiological metrics

As neuronal activity was lower in the presence of microglia, we sought to investigate whether neurons were still functional and able to respond to stimulation or whether this capacity was disrupted in presence of microglia. Increased neuronal activity in response to the potassium channel blocker 4-Aminopyridine (4-AP) has previously been characterised in other cell models (Parmentier et al., 2022). Neuronal activity in both cocultures and tricultures from three independent biological repeats was modulated with 4-AP (Fig 5). Treatment at DIV17 significantly increased the mean firing rate in cocultures and tricultures compared to their basal activity before drug treatment (Fig 5A, N/AS *p* = 0.0265, N/AS/MG^med^ *p* < 0.0001, N/AS/MG^high^ *p* < 0.0001). Treatment also increased burst frequency in cocultures and tricultures (Fig 5B, N/AS *p* = 0.0004, N/AS/MG^med^ *p* < 0.0001, N/AS/MG^high^ *p* < 0.0001). Network burst frequency was also increased by 4-AP treatment (Fig 5C, N/AS *p* = 0.0095, N/AS/MG^med^ *p* = 0.0068, N/AS/MG^high^ *p* = 0.001). The N/AS/MG^high^ triculture displayed a larger increase in mean firing rate and burst frequency compared to N/AS or N/AS/MG^med^, whereas the percentage increase in network burst frequency was more consistent between cocultures and tricultures. Representative activity traces (raster plots) at DIV17 from N/AS/MG^high^ tricultures before and after the 4-AP treatment show the increase in neuronal activity after drug addition (Fig 5D and 5E). In conclusion, modulation of cortical neuronal activity in cocultures and tricultures with 4-AP indicates that neurons in the triculture model remain functional and retained the capacity for neuronal activity in the presence of the microglia.

**Figure 5:**
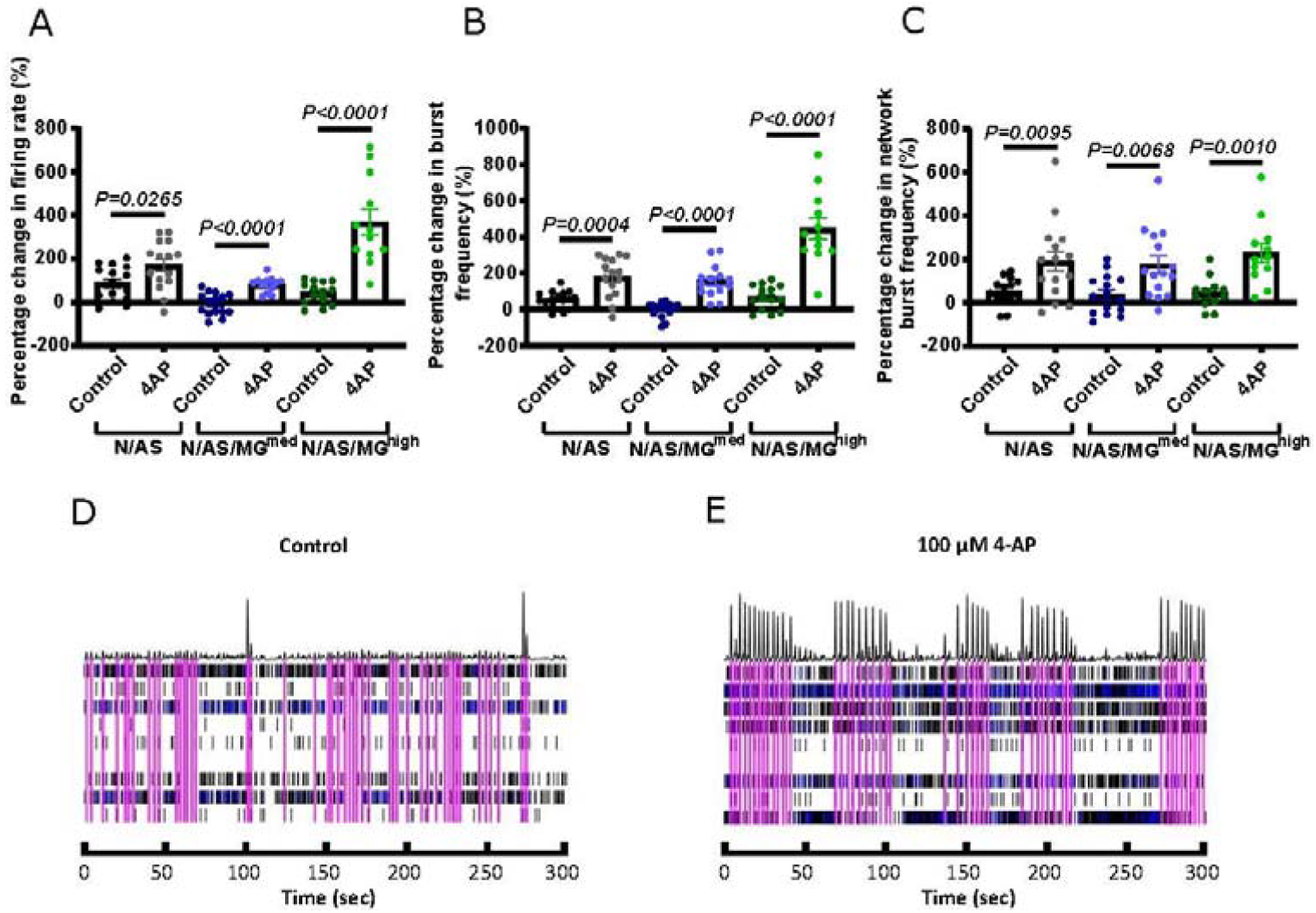
Treatment with 4-Aminopyridine (4-AP) increases cortical neuronal activity in cocultures and tricultures. (**A**). On DIV17, cultures were treated with vehicle or 100 μM 4-AP. Scatter plot shows the percentage change in mean firing rate of neurons 1 hour after treatment compared to their basal activity before drug treatment. (N/AS *p* = 0.0265, N/AS/MG^med^ *p* < 0.0001, N/AS/MG^high^ *p* < 0.0001, Unpaired t test). (**B**). On DIV17, cultures were treated with vehicle or 100 μM 4-AP. Scatter plot shows the percentage change in burst frequency of neurons 1 hour after treatment compared to their basal activity before drug treatment. (N/AS *p* = 0.0004, N/AS/MG^med^ *p* < 0.0001, N/AS/MG^high^ *p* < 0.0001, Unpaired t test). (**C**). On DIV17, cultures were treated with vehicle or 100 μM 4-AP. Scatter plot shows the percentage change in network burst frequency of neurons 1 hour after treatment compared to their basal activity before drug treatment. (N/AS *p* = 0.0095, N/AS/MG^med^ *p* = 0.0068, N/AS/MG^high^ *p* = 0.001, Unpaired t test). (**D, E**). Representative activity traces at DIV17 from triculture N/AS/MG^high^ treated with vehicle and 100 μM 4-AP. Black spikes at the top represent network activity while activity over time recorded on each of the 8 electrodes is shown under. Blue = bursts. Pink = network bursts. Data shown as mean ± SEM and is representative of 3 independent experiments.

### 2.4. Microglia reduce the densities of spines and the postsynaptic density scaffold protein Homer1 in secondary dendrites

Following our finding that microglia suppress spontaneous neuronal activity, we further investigated the impact of microglia on neurons in the triculture model. Neuronal cultures, cocultures and tricultures were transfected with Synapsin-GFP to visualise individual dendrites and spines before immunostaining for the presynaptic protein VGlut1 and postsynaptic protein Homer1 (Fig 6A-E). Dendritic spine density is reduced in neuronal cultures and tricultures compared to cocultures. The trend indicates that microglia reduce spine density in a dose-dependent manner, with N/AS/MG^high^ tricultures displaying similar spine densities to that of the neuronal cultures. The difference in spine densities between the five culture conditions is statistically significant (*p* = 0.0263) (Fig 6F). While not significant, the density of Homer1 puncta in neuronal cultures, cocultures and tricultures follows a similar trend, albeit with more variation between biological repeats (*p* = 0.0839) (Fig 6G). The percentage of Homer1 puncta colocalised with VGlut1 does not indicate a clear trend (data not shown).

**Figure 6:**
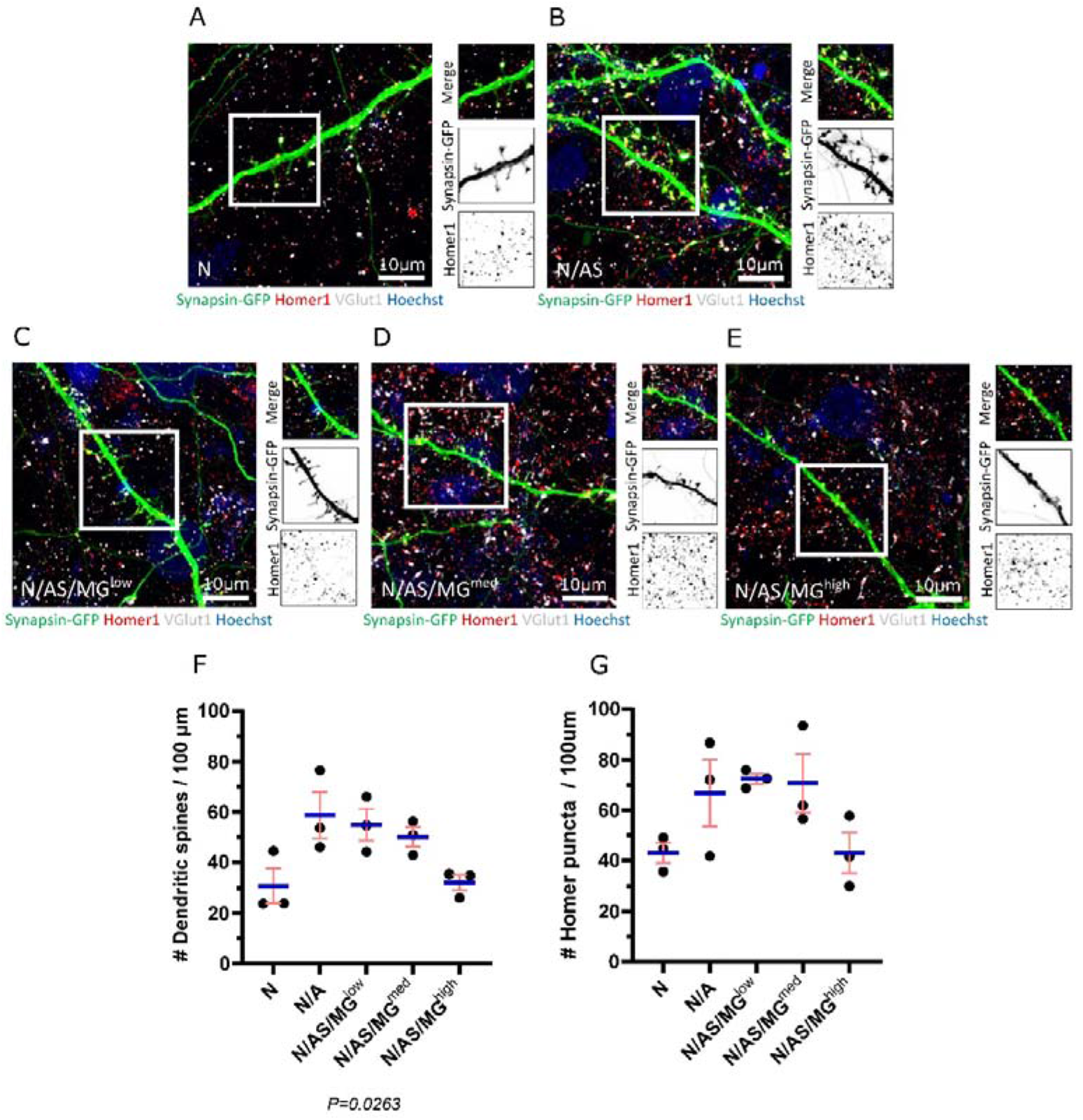
Spine density and Homer1 density are reduced in neuronal cultures and tricultures compared to cocultures. (**A-E**). Representative immunostaining images of dendrites used for quantification of spines and Homer1 density showing Synapsin-GFP, Homer1 and VGlut1 expression at DIV17 in neuronal cultures, cocultures and tricultures. Image shows staining of Synapsin-GFP (green), Homer1 (red), VGlut1 (white) and Hoechst (blue). Scale bar = 10 μm. White frame = field of view of Merge. (**F, G**). Analysis performed on neuronal cultures, cocultures and tricultures to quantify the number of spines and Homer1 puncta using IMARIS software. The difference in spine densities between the five culture conditions is statistically significant (*p* = 0.0263, one-way ANOVA). The difference in Homer1 densities between the five culture conditions is not statistically significant (*p* = 0.0839, one-way ANOVA). Data shown as mean ± SEM and is representative of 3 independent experiments.

## 3. Discussion

We developed an *in vitro* rodent triculture model of neurons, astrocytes and microglia that is suitable for pharmacological modulation. The triculture is established by adding exogenous microglia to cocultures of neurons and astrocytes and culturing the cells in a triculture medium supplemented with TGF-β, IL-34 and cholesterol. These three components were chosen as they are the key factors sufficient for promoting microglial survival under serum-free conditions (Bohlen et al., 2017) and have been used previously to generate a primary triculture model (Goshi et al., 2020). Immunostaining shows that there is a healthy population of neurons and astrocytes in both the cocultures and tricultures, and the number of neurons is not affected by the presence of the microglia. The number of astrocytes appears to be slightly reduced in the triculture with the highest ratio of microglia. While it is possible that only expression of Sox9 is affected, lower numbers of astrocytes have previously been observed in tricultures compared to cocultures by quantifying the number of nuclei co-localised with glial fibrillary acidic protein (GFAP) (Goshi et al., 2020). One possible explanation is that microglia express TGF-β1, which has been shown to inhibit astrocyte proliferation (Lindholm et al., 1992; De Groot et al., 1999). Further analysis indicates that the percentage of homeostatic microglia decreases dose-dependently as the number of microglia increases. Therefore, tricultures with higher numbers of microglia contain a microglia population that is proportionally more reactive.

By using MEA as an endpoint to investigate glia-mediated regulation of cortical neuronal activity, we have demonstrated that neuronal cultures exhibit lower basal activity than cocultures of neurons and astrocytes, which is consistent with the reduced spine and Homer1 densities displayed in our neuronal cultures. Previous studies have also shown that astrocytes increase the number of mature, functional synapses on neurons (Ullian et al., 2001). This is thought to occur via the secretion of factors such as thrombospondins which are known to promote synaptogenesis (Christopherson et al., 2005). We also demonstrated that tricultures exhibit lower basal activity than cocultures and that microglia suppress neuronal activity in a dose-dependent manner. Pharmacological modulation of these cultures with 4-AP indicates that neurons remain functional in the presence of the microglia even if their basal activity is lower. Microglia have previously been shown to regulate neuronal activity. Neurons generate ATP, which attracts microglial processes to neurons with high levels of activity. In zebrafish this microglia-neuron contact requires the small Rho GTPase Rac in microglia, and results in reduced activity in the contacted neurons (Li et al., 2013). Microglia also respond to neuronal activation by suppressing neuronal firing and the associated seizure response in mice. The mechanism of suppression is dependent on the microglial sensing of ATP and its subsequent conversion into adenosine (Badimon et al., 2020). Another possible mechanism for the downregulation of neuronal activity in the triculture model is complement-mediated synapse elimination. This hypothesis is supported by the fact that microglia reduce the densities of spines and Homer1 puncta in a dosedependent manner in our triculture model. The formation of mature neural circuits requires activity-dependent pruning of inappropriate, less active synapses, and this process occurs via activation of C3 receptors on microglia, triggering phagocytosis (Stevens et al., 2007). The complement-dependent synaptic pruning that occurs during development is thought to be reactivated and contribute to synapse loss in AD (Hong et al., 2016). Microglia may also reduce synapse numbers via a non-phagocytic mechanism. In the retinogeniculate circuit, microglia expressing TNF-related weak inducer of apoptosis (TWEAK) facilitate synapse elimination through the neuronal receptor Fibroblast growth factor-inducible 14 (Fn14) (Cheadle et al., 2020).

It is important to note that while our model can provide insights into the cross-talk between microglia, astrocytes and neurons, it may not fully recapitulate the interactions involved in human disease. Cultures generated from rodents are an important research tool and have contributed heavily to our understanding of disease mechanisms. In many respects they mirror human biology closely, and the genes identified as being associated with inherited disease in humans are highly conserved in the rat genome (Huang et al., 2004). However, understanding species-specific differences in the neuroinflammatory response remains a major challenge in the field and there are recognised limitations in using mouse models to mimic human disease, specifically neurodegeneration (Arber et al., 2017). Further development of our model could incorporate cell types of other species. A human triculture system of hiPSC-derived microglia, astrocytes and neurons containing the APPSWE+/+ mutation has been used to model AD and showed increased production of the complement C3 protein due to microglia initiating reciprocal signalling with astrocytes (Guttikonda et al., 2021). While iPSCs provide a valuable tool for disease modelling using physiologically relevant cells, they can prove challenging when attempting to generate a robust, reproducible culturing platform for drug discovery. Automated technology has been utilised to eliminate the variation that can arise during differentiation and produce consistent and long-term tricultures of hiPSC neurons, astrocytes and microglia. This model displays many pathological features of AD such as Aβ plaques, phospho-Tau induction and synapse loss, and has been used to investigate the mechanism of action of anti-Aβ antibodies (Bassil et al., 2021). Mixed species cultures provide additional opportunities to probe the roles of individual cell types within a more complex model. RNA-sequencing on cultures combining rat, mouse and human cells allows for individual cell-type transcriptomes to be profiled. This approach has been used to show that microglial ramification is controlled by the combined effects of neurons and astrocytes via TGF-β2 signalling (Baxter et al., 2021).

Most AD drug targets which displayed favourable outcomes in biochemical, cell culture and AD transgenic models have failed to prove effective in clinical trials (De Strooper, 2014). One possible explanation for this is the limited ability of these models to mimic human disease. In 2D cell culture models of AD, secreted Aβ may diffuse into the cell culture media and be removed during media changes (D’Avanzo et al., 2015). Recent advances in stem cell and 3D culture technologies have made it possible to generate novel models that more closely recapitulate AD pathology in the human brain. A neural stem-cell-derived 3D culture system generated using Matrigel-based technology behaves more like brain tissue and provides a closed environment promoting aggregation of Aβ. In this model, familial AD mutations in *β-amyloid precursor protein* and *presenilin 1* induce robust extracellular deposition of Aβ, including Aβ plaques, and aggregates of phosphorylated tau in the soma and neurites (Choi et al., 2014). While these 3D cell culture models provide a promising, more physiologically relevant platform for high-throughput drug screening, neurospheroids can be sensitive to small variations and difficult to manipulate experimentally. Recently however, a platform has been developed using new tools and stem cell engineering to reproducibly generate neurospheroids inside a 96-well cell culture array plate with 1,536 microwells. Unlike other brain organoids, in this model the dendrites extend outward which allows for the formation of networks (Jorfi et al., 2018).

As neuron-glia interactions become increasingly recognised to play a critical role in the pathogenesis of AD, the development of complex models and endpoints allowing for modulation of glia within a system has become more important for drug discovery. While 3D models allow for more complex modelling of disease, the 2D model we describe here may provide a more realistic alternative for many studies as it is easier to generate and can be used to measure physiologically relevant endpoints. Numerous studies have targeted glia in models of neurodegenerative diseases. For example, pharmacological blockade of Connexin 43 (Cx43) in ALS astrocytes has been shown to provide neuroprotection of motor neurons and reduce neuronal hyperexcitability in a coculture model (Almad et al., 2022). Inhibition of astrocytic α2-Na+/K+ adenosine triphosphatase (α2-NKA) in a mouse model of tauopathy suppresses neuroinflammation, and its knockdown halts the progression of tau pathology (Mann et al., 2022). Blocking of the astrocytic calcium channel Transient Receptor Potential Ankyrin 1 (TRPA1) or the enzyme epoxide hydrolase, which is predominantly expressed by astrocytes, have both been shown to normalise astrocytic activity, prevent neuronal dysfunction and improve cognitive function in AD mouse models (Ghosh et al., 2020; Paumier et al., 2022). TREM2 is essential for the transition of homeostatic microglia to a disease-associated state. Monoclonal antibody-mediated stabilisation and activation of TREM2 on the cell surface reduces AD-related pathology in a mouse model and reduces levels of the homeostatic marker P2RY12, suggesting that driving microglia towards a disease-associated state might provide a protective function (Schlepckow et al., 2020). Activation of TREM2 in an AD mouse model modulates the microglial inflammatory response, reduces neurite dystrophy and restores the behavioural changes associated with Aβ pathology (Wang et al., 2020).

In summary, our neuron, astrocyte and microglia triculture model provides a robust and reliable tool for studying the role of glia-neuron crosstalk in the regulation of neuronal activity, and how this is impacted by neuroinflammatory processes in disease. The use of this model in combination with MEA technology allows for pharmacological manipulation of the system in a high-throughput manner and has the potential to be used for target validation and drug screening.

## 4. Experimental procedures

### Animals

All procedures involving animals were conducted according to the UCL Ethics Committee guidelines and the UK Animals Scientific Procedures Act UK (1986) and its Amendment Regulations (2012). Sprague-Dawley rats (Charles River) were used in this study.

### Primary cortical neuron and astrocyte cocultures

Cocultures were prepared from the neocortex of embryonic day 18 (E18) embryos of Sprague-Dawley rats. Meninges were removed and the cerebral cortices were dissected. The tissue was then incubated with 0.25% trypsin-EDTA (Gibco, 25200056) and DNase I (50 μg/ml, Sigma-Aldrich) for 10 min at 37°C. The cells were mechanically dissociated in DMEM (Gibco, 31966021) complemented with DNase I and washed twice in DMEM. 25000 neurons/well were plated on a 96-well imaging plate coated overnight with Poly-D-Lysine (PDL) (Sigma, P0899), and 50000 neurons/well were plated on a 96-well MEA plate (see MEA Methods for detailed protocol). The cultures were maintained for 24 hours in a plating media composed of DMEM supplemented with Glutamax (Gibco, 35050061), 5% horse serum (HBS) (Gibco, 26050088), 1mM glutamine (Gibco, 25030081) and penicillin/streptomycin (Gibco, 15140148). After one day in vitro (DIV1), medium was removed and replaced with maintenance media composed of Neurobasal (Gibco, 21103049), B27 1x (ThermoFisher, 17504044), N2 1x (ThermoFisher, 17502048), 1mM glutamine (Gibco, 25030081), Glutamax (Gibco, 35050061) and penicillin/streptomycin (Gibco, 15140148). Twice a week, half of the medium was removed and replaced with fresh medium. The cultures were maintained in a humidified incubator at 37°C under an atmosphere containing 5% CO2. Cocultures were used for experimentation between DIV13 and DIV17.

### Primary cortical neuronal cultures

Neuronal cultures were initially prepared as above, using the same method as the cocultures. Cultures were plated for 4 hours in plating media. After 4 hours, medium was removed and replaced with fresh maintenance media. Twice a week, half of the medium was removed and replaced with fresh medium. On DIV6, medium was supplemented with CultureOne™ Supplement (Gibco, A3320201). The cultures were maintained in a humidified incubator at 37°C under an atmosphere containing 5% CO2.

### Primary microglia culture

Microglia were prepared from postnatal day 14 (P14) wildtype Sprague-Dawley rats. The meninges, cerebellum and olfactory bulbs were removed before the hemispheres were minced into small pieces using a sharp blade. Minced tissue was mechanically dissociated in a glass homogeniser in PBS, centrifuged at 1380g for 10 minutes and the pellet resuspended in 70% isotonic Percoll (Sigma, P1644). Layers of 30% isotonic Percoll and PBS were slowly added before centrifuging at 1090g for 50 minutes with slow acceleration/deceleration. The myelin layer was discarded from the top of the Percoll gradient, and the microglia collected from the interface between the 70% and 30% Percoll layers. Microglia were diluted in PBS and centrifuged at 1570g for 10 minutes and the pellet resuspended in DMEM/F12 containing penicillin/streptomycin (Gibco, 15140148), 2mM L-glutamine (Gibco, 25030024), 5 μg/ml N-acetyl cysteine (Sigma, A8199), 5 μg/ml insulin (Sigma, I6634), 100 μg/mL apo-transferrin (Sigma, T1147), and 100 ng/mL sodium selenite (Sigma, S5261) (basal media) supplemented with 2 ng/mL TGF-β2 (Peprotech, 100-35B), 100 ng/mL IL-34 (BioLegend, 577606), 1.5 μg/mL cholesterol (Avanti, 57-88-5) and 1 μg/mL heparan sulfate (Amsbio, AMS.GAG-HS01) (TICH media). Cultures were maintained in a humidified incubator at 37°C under an atmosphere containing 5% CO2.

### Primary neuron – astrocyte – microglia triculture

Cocultures were prepared as above and cultured in maintenance media until DIV6, from which point cells were cultured in maintenance media supplemented with TGF-β2 (2 ng/mL), IL-34 (100 ng/mL) and cholesterol (1.5 μg/mL) (triculture media). On DIV8, microglia were isolated and centrifuged at 1570g for 7 minutes to remove the TICH media. The pellet was resuspended in triculture media and 20 uL of microglia was added to the coculture in ratios of 1 microglia:10 neurons/astrocytes, 1:3 and 1.5:1. Twice a week, half of the medium was removed and replaced with fresh medium. Cultures were maintained in a humidified incubator at 37°C under an atmosphere containing 5% CO2. Tricultures were used for experimentation between DIV13 and DIV17.

### Transfection

Neuronal transfection with the DNA construct pHR hsyn:EGFP (Keaveney et al., 2018; kind gift from Xue Han (Addgene plasmid #114215; http://n2t.net/addgene:114215; RRID:Addgene_114215)) was performed at DIV7 using the Neuromag magnetofection method (OzBiosciences). For each well of a 96-well plate, 1.5 μg DNA was diluted in 100 μl of Opti-MEM and added dropwise to 1 μl Neuromag transfection reagent, with all reagents at room temperature. Following 20 minutes incubation at room temperature, the transfection mix was added dropwise to neuronal cultures and the 96-well plate was placed on a magnetic plate (OzBiosciences) pre-equilibrated to 37 °C inside an incubator. After 15 minutes of magnetofection, the culture plate was removed from the magnetic plate and normal cell culture resumed.

### Immunocytochemistry

Cultures fixed in 4% PFA were labelled after membrane permeabilization and saturation with 10% FBS and 0.02% Triton X-100 (Sigma) in PBS for 1 hour. Cells plated for imaging were incubated with mouse NeuN (1:500, ab104224, Abcam), rabbit Sox9 (1:500, ab185230, Abcam), guinea pig Iba1 (1:500, 234 004, Synaptic Systems) and rabbit Olig2 (1:500, ab109186, Abcam). Cells plated in MEA plates were incubated with mouse NeuN (1:250, ab104224, Abcam), rabbit Sox9 (1:250, ab185230, Abcam) and guinea pig Iba1 (1:500, 234 004, Synaptic Systems). Antibodies were diluted in in 10% FBS and 0.02% Triton X-100 in PBS and incubated overnight at 4°C. After rinsing three times in PBS, cells were incubated with anti-mouse Alexa Fluor 488, anti-guinea pig Alexa Fluor 568 and anti-rabbit Alexa Fluor 647 diluted 1:1000 in PBS with 10% FBS and 0.02% Triton X-100 for 2 hours at room temperature. Nuclei were stained with Hoechst 33258 (1:1000, 94403, Sigma-Aldrich) and cells washed three times in PBS.

For synaptic staining, transfected cultures fixed in 4% PFA and 4% sucrose were labelled after membrane permeabilization and saturation with 5% NGS, 1% BSA and 0.1% Triton X-100 in PBS for 1 hour. Cells were incubated with rabbit Homer1 (1:500, 160 003, Synaptic Systems) and guinea pig VGlut1 (1:300, AB5905, Merck Millipore). Antibodies were diluted in 5% NGS, 1% BSA and 0.1% Triton X-100 in PBS and incubated overnight at 4°C. After rinsing three times in PBS, cells were incubated with anti-rabbit Alexa Fluor 568 and anti-guinea pig Alexa Fluor 647 diluted 1:500 in PBS with 5% NGS, 1% BSA and 0.1% Triton X-100 for 2 hours at room temperature. Nuclei were stained with Hoechst 33258 (1:1000, 94403, Sigma-Aldrich) and cells washed three times in PBS.

### Image acquisition

For cultures used for cell quantification, images were acquired using the Opera Phenix High-Content Screening System from PerkinElmer. Nine fields of view were taken at 10x magnification to scan an entire 96-well culture well. To analyse the morphology of microglia in the tricultures, images were again acquired using the Opera Phenix High-Content Screening System from PerkinElmer. Nine fields of view were taken at 20x magnification to obtain representative images spread across each 96-well culture well.

For cultures on MEA plates, images were acquired using the Celldiscoverer 7 Automated Microscope from Zeiss. Thirty six fields of view were taken at 5x magnification to scan an entire 96-well culture well.

For transfected cultures used for synaptic staining, images were acquired using the LSM880 Confocal from Zeiss. Whole neurons were acquired using a 40x objective (NA = 1.3), at a resolution of 1024 x 1024 pixels, a step size of 1 μm, and with photomultiplier tubes to detect fluorescence emission. Secondary dendrites were acquired at a higher magnification using the 63x objective (NA = 1.4) with an additional 3.5x zoom at a resolution of 1024 x 1024 pixels and at a step size of 0.5 μm. A total of 6 neurons and accompanying secondary dendrites spread across three biological repeats were imaged per condition.

### Image analysis

The Harmony High-Content Imaging and Analysis Software was used to quantify the neuron, astrocyte and microglia cell numbers in culture. Microglia were classified into either amoeboid or ramified phenotypes using the PhenoLOGIC machine-learning algorithm within the Harmony software, with modifications to the training of the algorithm to enable recognition of microglia within the tricultures.

Dendritic spine and synapse analysis on hSyn:EGFP expressing cortical neurons was performed using IMARIS software. Briefly, the ‘Filament’ tool was used to semi-automatically specify the secondary dendrite within an image file, followed by the detection of dendritic spines by manual identification. The GFP signal was used to create an exclusion mask using the ‘Surface’ tool to isolate the Homer1 signal within a dendrite of interest. Homer1 puncta were subsequently identified using the ‘Spot’ detection tool set to a detection diameter of 0.45 μm. Background Homer signal was excluded by thresholding using the ‘Quality’ filter.

Pre-synaptic vGlut1 puncta in the entire image were similarly identified using the ‘Spot’ detection tool at 0.45 μm diameter. Synapses were assumed using the ‘Colocalize spots’ function within the ‘Spot’ detection tool when there was a maximum distance of 1 μm between Homer1 and vGlut1 puncta.

### Multi-Electrode Array (MEA)

Each well of a 96-well MEA plate (M768-tMEA-96B, Axion Biosystems) was spot-coated with 8 μL of PEI solution (0.2% PEI solution in 0.1M sodium borate pH 8.4) and plates were incubated at room temperature overnight. Wells were washed three times with water before being left to dry under the biological safety cabinet for approximately one hour. Wells were then spot-coated with 8uL of laminin (20μg/mL; Sigma L2020). Sterile deionized water was added to the area surrounding the wells to prevent substrate evaporation and the plates were incubated for 1 hour at 37C without letting the laminin droplet to dry. Laminin was removed directly before seeding the well with neurons. 50000 neurons/well were spot-coated in 8uL of plating media. 10 replicates were plated per condition in the inner wells of the plate. Plates were incubated for 1h at 37C before 150uL of plating media was added to each well. On the next day cells were transferred in maintenance media and maintained as described above. Electrical activity of neurons was recorded daily between DIV13 and DIV17. When recordings took place on the same day as the half-media changes, neurons were first recorded as media changes will temporarily alter neuronal activity. On the recording day, the plate was loaded into the Axion Maestro MEA reader (Axion Biosystems), let to equilibrate for 10 minutes and recording was performed via AxIS for 15 minutes (1000 X Gain, 200 Hz – 3 kHz). Data was first analysed with AxIS software (Spike Detector Threshold 6 x Std Dev) and then subsequently using GraphPad Prism 9 software.

### Drug treatment

On DIV17, half an hour after basal neuronal activity was recorded, the MEA plate was treated by removing half of the medium and replacing with medium containing 100 μM 4-Aminopyridine (275875, Sigma-Aldrich) at 2X. Recordings were performed 1 hour after treatment.

For analysis, wells were excluded using the ROUT method to detect outliers (Q=10%) in GraphPad Prism 9 (GraphPad Software, CA, USA).

### Statistical analysis

All mean values are presented and stated as ± standard error of mean (SEM). Statistical analyses were performed using GraphPad Prism 9 (GraphPad Software, CA, USA). Data normality was tested by the Shapiro-Wilk test.

Comparisons between two groups were analysed by unpaired t test. One-way ANOVA was used for comparisons between >2 groups in datasets with one variable. 2-way ANOVA was used for comparisons between >2 groups in datasets with two variables. Post-hoc Tukey’s multiple comparisons test was used after performing a one- or 2-way ANOVA.

## Acknowledgements

This work was supported by Alzheimer’s Research UK (grants ARUK-2018DDI-UCL and ARUK-2021DDI-UCL).

